# dirHub: a trackHub configurator with directory structure projection

**DOI:** 10.1101/314807

**Authors:** Hideya Kawaji

## Abstract

**Summary:** Track Data Hub is a mechanism enabling us to visualize genomics data as tracks along genome coordinates and share them over the Internet, relying on a web server hosting data files and genome browsers offering graphical representations. It requires an accessible configuration file specifying all graphical parameters and track hierarchy, in addition to the data files. Here dirHub is developed to assist generation of the configuration file by projection of a file directory structure, which makes it possible to set up trackHub visualization mostly by file operations.

**Availability and implementation:** It is implemented in ruby and the source code is available at https://github.com/hkawaji/dirHub/. It is tested on the UCSC Genome Browser and the Hub Track Database Definition (v2).

## Introduction

With the progress of sequencing technologies, a large amount of genome wide data, such as epigenetic and transcriptome profiles as well as genome sequences, have been being produced. It is crucial to obtain integrated perspective on such data, including newly produced data as well as datasets already available in public. One of the most basic processes to understand such genomics data is its graphical representation as a “track”, where the contents, such as monitored events or signals, are schimaically represented along genome coordinates.

trackHub (Raney *et al.*, 2014) provides an efficient scheme to generate and share track visualizations, where a user makes his/her own data files available on the internet and a server of genome browser access them to make their graphical representation. The scheme is supported by major genome browsers such as the UCSC genome browser database (Casper *et al.*, 2018), ENSEMBLE genome browser (Zerbino *et al.*, 2018), Dalliance genome browser (Down *et al.*, 2011), and it is also extensively employed by large collaborative efforts such as ENCODE (Sloan *et al.*, 2016), Roadmap epigenomics (Karnik and Meissner, 2013), blueprint (http://dcc.blueprint-epigenome.eu/), and FANTOM5 (Lizio *et al.*, 2015). It is valuable not only for such large consortia to distribute their results, but also for individual studies to have an integrate view of their own datasets with other related data sets.

Our goal here is to facilitate trackHub-based visualization by assistance of its configuration. A set of data files, typically in bam (Li *et al.*, 2009) or bigBed/bigWig (Kent *et al.*, 2010) format, are produced through a series of data processing steps. Graphical parameters described in the specification (https://genome.ucsc.edu/goldenpath/help/trackDb/trackDbHub.html) are described in a configuration file to visualize each of the data files as a track. Grouping of multiple data can be achieved by setting a container of multiple tracks, such as multiWig, composite, and super track, which is suitable in particular to organize a large set of data files into meaningful chunks. All these settings have to be described in a configuration file, which typically require a small script or program made only for the data set. The tool we developed here, dirHub, enables us to generate a trackHub configuration file by mapping a data file to a single track according to suffix of its file name, and a directory to a track container. Since it is owing to the projection of a file directory structure to a track hierarchy, trackHub-based visualization can be configured mostly by file operations.

## Results

### Mapping data file structure to track type

Conventionally suffix of a file name is used to specify a data type, and we follow such conventions when available. For example, “.bb” in the end of the file name indicates bigBed file (Kent *et al.*, 2010). However there is no conventional suffix for “container” type track since there is no corresponding data file. Here we use a file directory to represent such a container track. Suffixes “.mw”, “.cp”, and “.st” of directory name are used to indicate a “multiWig”, “composite track”, and “superTrack” type container, and data files in the directory are considered as tracks included in the container (Fig 1a,b).

**Figure 1.**
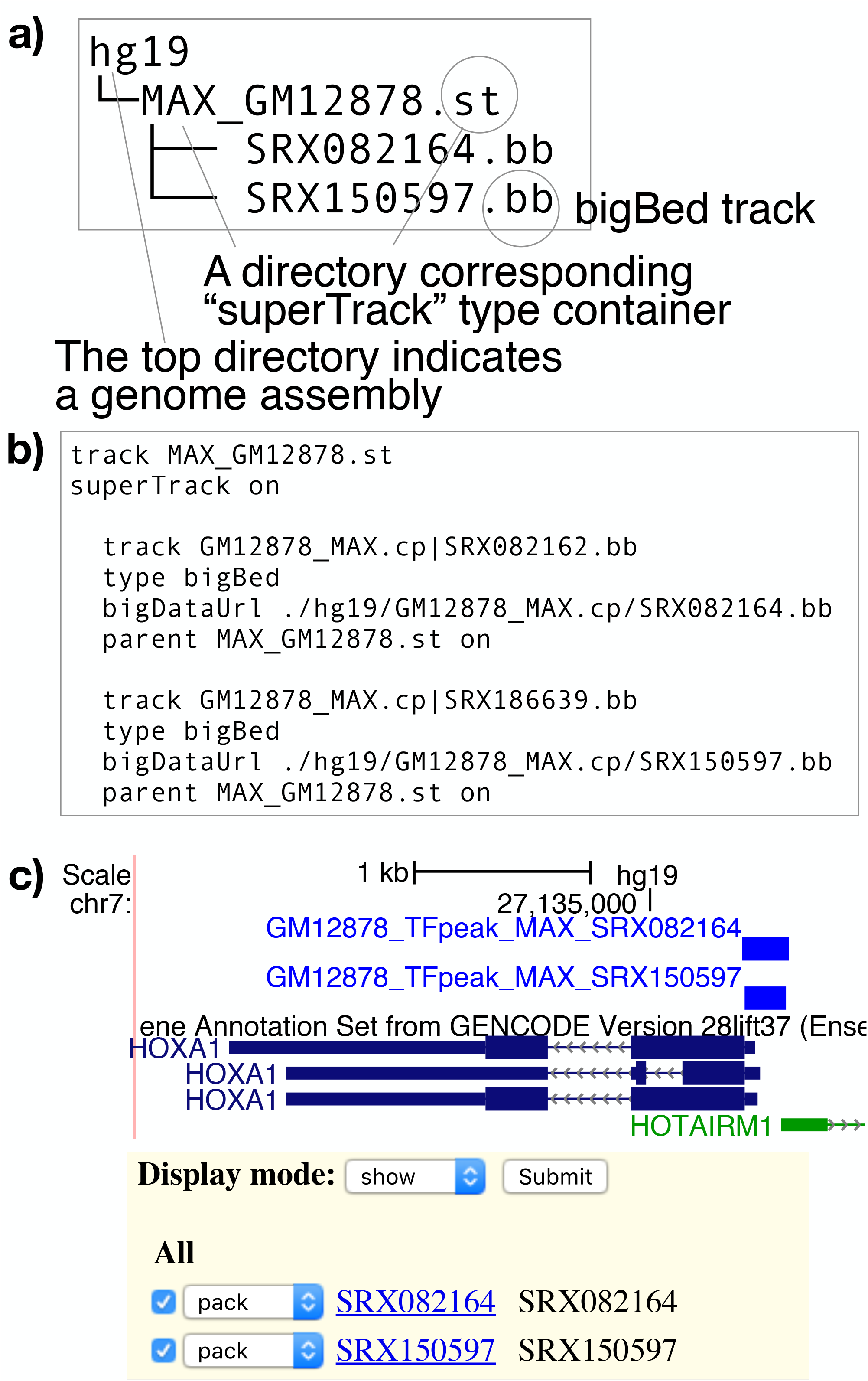
An example of data file directory and its projection to a trackHub configuration. (a) A data file directory consisting of two bigBed files. The “hg19” directory is prepared to indicate the genome assembly, on which the tracks are based on. Suffixes of “.st” and “.bb” indicates track types, “superTrack” and “bigBed” respectively. The two bigBed tracks belong to a superTrack container. (b) A configuration file that is projected from the directory structure. Mandatory attributes, shortLabel and longLabel were omitted here for explanation. (c) A graphical representation of these tracks with its user interface.

All tracks have to be tied to a genome assembly to specify the primary axis where the graphical representation should be based on. We require the top of data file structure to have a directory which name is identical to the assembly name (Fig 1a). By using a configuration file generated by directory structure projection, trackHub can be set up as in Fig1c.

### Two ways to generate trackhub configuration

dirHub can be used in two ways, as a web service, or a command line tool (Fig 2). In case of using its web interface (Fig 2b), the dirHub server host the generated configuration file on the internet, which can be immediately subjected to use of trackHub (that is, the URL of the configuration file can be pasted in “myHub” box). It has to be noted the response in trackHub browsing could be relatively slow, since the configuration file is generated per request on the fly. It would be suitable for optimizing step of trackHub configuration, which requires many iterations of try and error, rather than intensive inspection of the data set after optimization.

**Figure 2.**
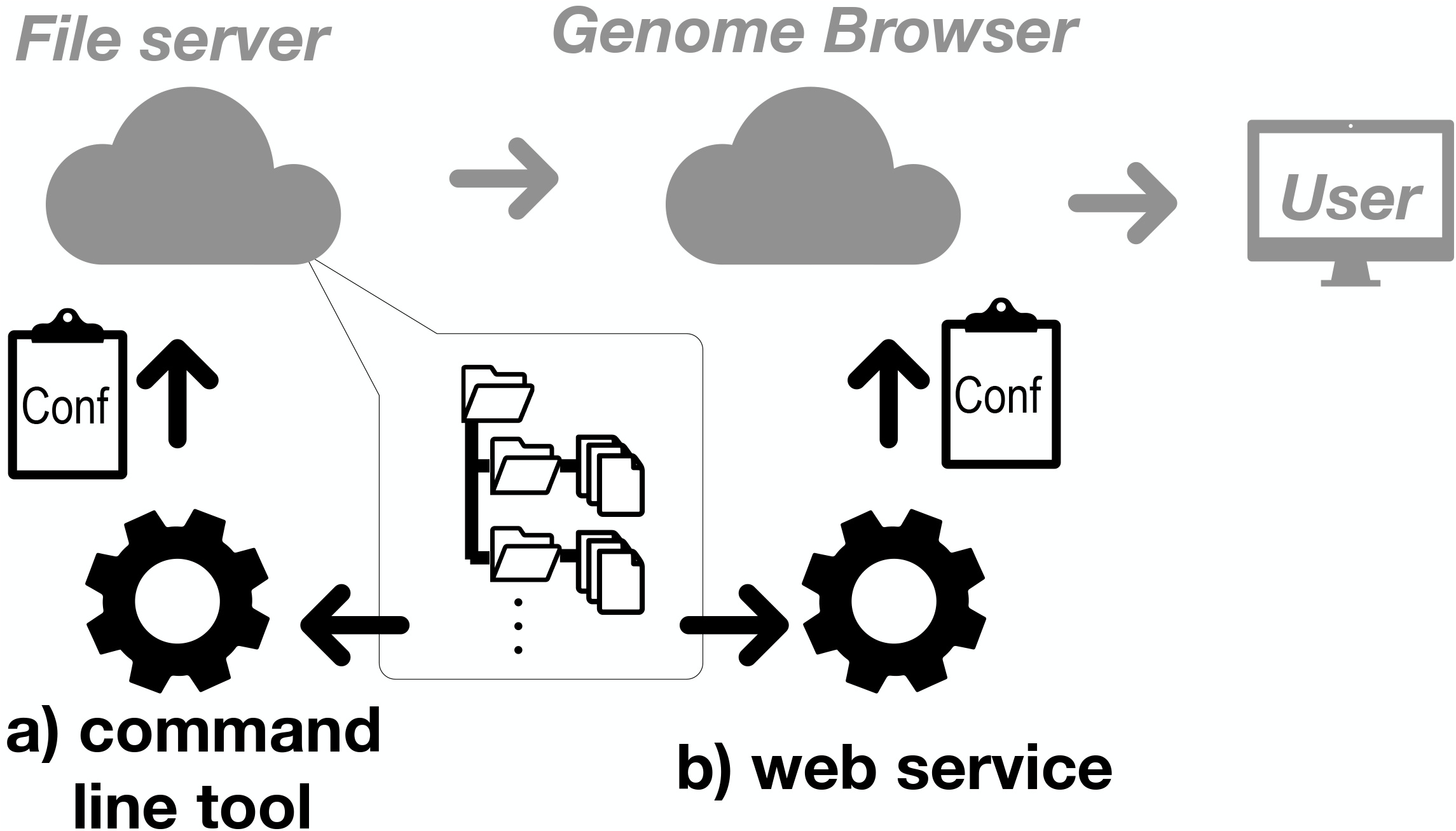
Two interfaces of dirHub. It can be used as a command line tool, which traverses the directory structure on the file server and generate configuration file there (a). Otherwise, it can be used as a web server, which traverses the directory structure over the internet and serve the generated configuration file to a genome browser by itself.

It also has to be noted that dirHub server intrinsically requires that contents of the data directory is visible on the internet (http/https). Listing the directory contents can be set up for example by using “DirectoryIndex” directive in apache http server (http://httpd.apache.org/docs/trunk/mod/mod_dir.html).

In the case of using its command line interface (Fig 2a), dirHub generates the configuration file on the local file system. It is applicable even to the case of using a file server that does not support directory listing, or the case of intensive inspection require rapid responses. Since the produced configuration file remains on the file system, users can use it as a template for further parameter optimization.

### Use case

We introduce an example, which focuses on K562 and GM12878 cell lines, to demonstrate how durHub can be effectively used. Their epigenetic profiles have been extensively monitored by the ENCODE project (The ENCODE project consortium, 2012), and their transcription initiation activities were monitored in the highest resolution by the FANTOM5 project (Forrest *et al.*, 2014). Here we are interested in an integrated view of these genomics data as in Fig 3a.

**Figure 3.**
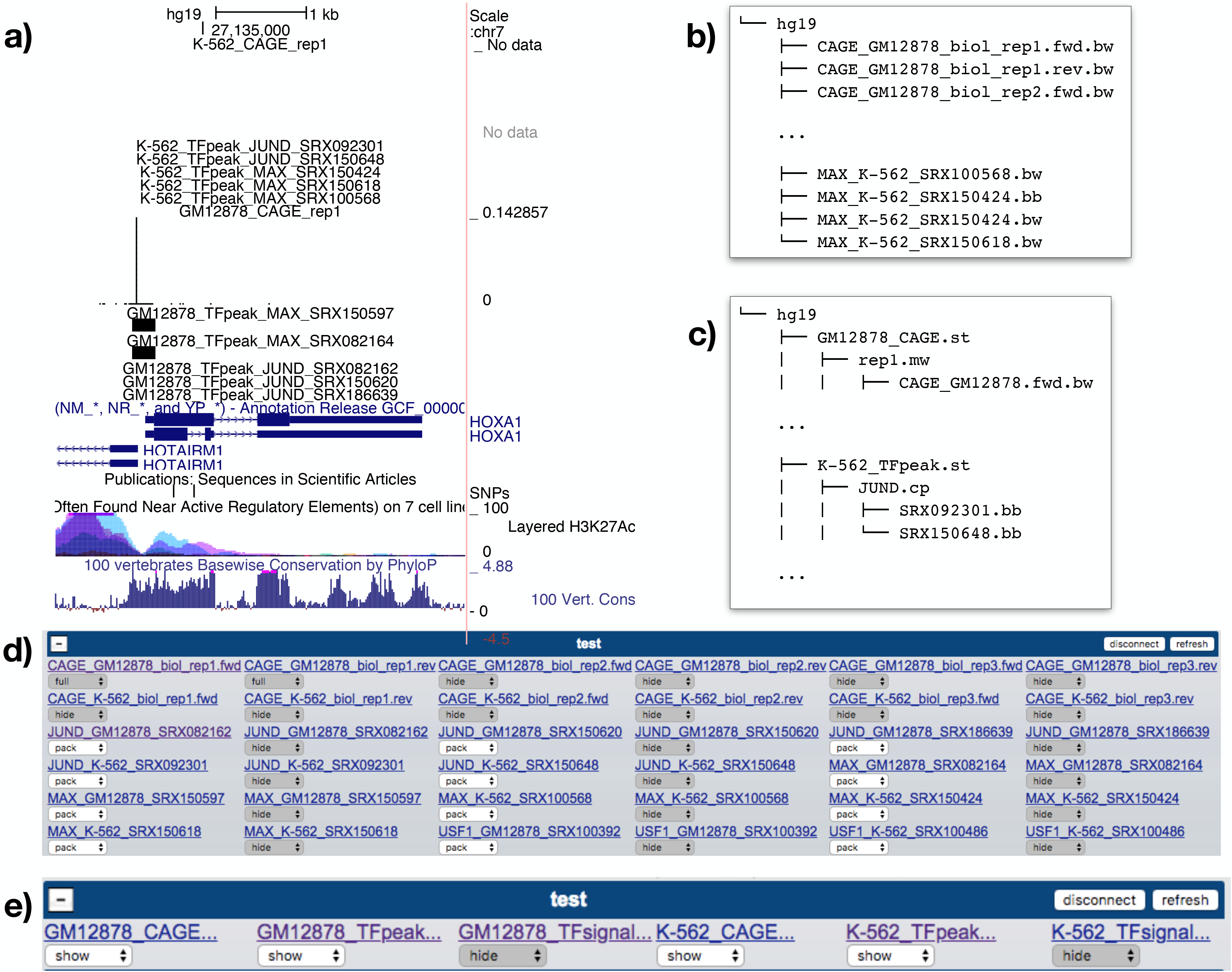
A use case of GM12878 and K562 profiling. Transcription initiation activities monitored by CAGE (Cap Analysis of Gene Expression) and epigenetic profiles of transcription factor (JUND and MAX) binding site monitored by ChIP-seq are displayed as tracks (a). A directory structure where all retrieved data files placed under the directory “hg19” (b), or structured depending on data types (c). The directory (b) and (c) are projected to trackHub configuration as in (d) and (e) respectively.

bigBed and bigWig data files of their epigenetic and transcriptome profiles were retrieved from ChIP-Atlas (Oki *et al.*, 2018) and the FANTOM5 web resource under a directory with a name of genome assembly, hg19 (Fig 3b). By giving URL of the directory containing the “hg19” directory to dirHub web interface, a configuration file is generated (Fig 3b). The URL can immediately be used in a genome browser to obtain track visualization (Fig 3d).

The process above provides an integrated view, but all data files are handled at the same level. It is not very straightforward to make operations on the view (such as turn on and off some tracks) on the interface. To group data files into cell lines and experiment types, the data files were reorganized as in Fig 3c. By projection of the directory structure, we can obtain a configuration which track containers were effectively used (Fig 3e). It has to be noted that all these processes relies in mostly on file operation.

## Summary

We introduced a framework to set up trackHub-based visualization. Its configuration file is generated by projection of the directory structure, trackHub-based visualization can be made mostly by file operations. It will facilitate rapid setup of trackHub and its iterative improvements.

## Material methods

dirHub is written in ruby, and tested with ruby version 2.3 and teackHub definition ver2. The source code is available at http://github.com/hkawaji/dirHub

## Acknowledgement

This work was supported by KAKENHI grants 16H02902, 16K12529, and National Bioscience Database Center (NBDC) of the Japan Science and Technology Agency (JST).

